# Ascl3+ ionocytes in murine salivary gland ducts are innervated sensory cells that display unique calcium signaling characteristics and contribute to the composition of saliva

**DOI:** 10.1101/2025.08.28.672671

**Authors:** Hitoshi Uchida, Eri O. Maruyama, Takahiro Takano, Marit H. Aure, Melissa F. Glasner, Zamira Guerra Soares, Roberta Faustoferri, V. Kaye Thomas, Helen P. Makarenkova, David I. Yule, Catherine E. Ovitt

**Author notes:** Hitoshi Uchida: Department of Oral Chrono-Physiology, Graduate School of Biomedical Sciences, Nagasaki University, 1-7-1 Sakamoto, Nagasaki, Nagasaki 852-8588 Japan. Eri O. Maruyama: The ADA Forsyth Institute, Somerville, MA 02143, USA. Marit H. Aure: National Institute of Dental and Craniofacial Research, National Institutes of Health, Bethesda, MD, USA. Melissa Glasner: The France Foundation, The Chandler Building, 84 Lyme Street, Old Lyme, CT 06371.

## Abstract

Ionocytes are distinct epithelial sensory cells scattered throughout the ductal system of the salivary glands. These cells are distinguished by their unusual morphology, as well as by a specific transcriptomic signature that includes expression of the Foxi1 and Ascl3 transcription factors. Currently, little is known about the biology or function of ionocytes in the salivary glands. To facilitate characterization of these cells, we generated an inducible Cre mouse allele driven by the *Ascl3* promoter. This strain was crossed with a reporter to fluorescently label Ascl3+ ionocytes, highlighting that they are the site of enriched CFTR expression in the salivary glands, and demonstrating the proximity of these cells to neurons. Conditional Cre-mediated cell ablation, using diphtheria toxin (DTA), removed Ascl3+ ionocytes from the salivary glands and resulted in an altered pH of total saliva, supporting a function for ionocytes in transepithelial ion flow. Finally, Cre-mediated expression of the calcium indicator GCaMP6f revealed that Ascl3+ ionocytes exhibit unique properties not observed in acinar or surrounding duct cells, including elevated basal [Ca^2+^]_I_, spontaneous blinking in the absence of stimulation, and a rapid loss of [Ca^2+^]_I_ following nerve stimulation. These unique properties distinguish ionocytes as a specialized subset of salivary gland duct cells.

## Introduction

Saliva is essential to maintain oral health. High concentrations of calcium and phosphate in saliva maintain tooth mineralization, high molecular weight proteins increase viscosity and prevent desiccation, while other proteins provide immune defense through direct anti-microbial action as well as promoting agglutination. In both mouse and humans, saliva is produced by three pairs of major salivary glands, the submandibular (SMG), sublingual and parotid. The majority of cells in each gland are clustered into acini, which secrete primary saliva into small intercalated ducts that are connected to an expanding ductal tree. While transporting the saliva to the oral cavity, cells in the duct system modify the isotonic fluid through mechanisms such as NaCl reabsorption, yielding the hypotonic total saliva. The intra- and inter-lobular segments of the salivary gland duct system vary between species, and are composed of a heterogeneous cell population, including innervated tuft cells, calcium mobilizing cells, and cells producing neurotrophic factors (Sato and Miyoshi, 1997, 1998; Shitara et al., 2007; Xiao et al., 2014; Yamamoto-Hino et al., 1998). The functional role of many of these duct cells remains poorly characterized.

Ionocytes were first identified in marine teleosts, where they are involved in the ionic balance of fluid secretion (Hwang et al., 2011). Recently, a novel sensory cell known as the ionocyte was described in pulmonary epithelia. This rare epithelial cell type comprises approximately 1% of lung epithelial cells (Montoro et al., 2018; Plasschaert et al., 2018). Identified through single-cell RNA Seq analysis, ionocytes have a unique transcriptional signature that includes the transcription factors forkhead boxI1 (Foxi1) and achaete scute-like 3 (Ascl3), multiple subunits of the vacuolar-type H(+)-ATPase (V-ATPase) and the cystic fibrosis transmembrane conductance regulator (Cftr), the gene that is mutated in cystic fibrosis. Pulmonary ionocytes are hypothesized to function as critical sensory cells in the airway epithelium (Xu et al., 2020), where they play a role in airway surface liquid absorption, secretion, pH and mucus viscosity (Lei et al., 2023; Yuan et al., 2023). Subsequent studies established that ionocytes are present in several tissues, including both human and murine salivary glands (Hauser et al., 2020; Huang et al., 2021; Mauduit et al., 2022). It is expected that they play a role in the modification of saliva, but the function of ionocytes in the salivary gland has not yet been established.

As in the lung, salivary gland ionocytes are marked by exclusive expression of Foxi1, and Ascl3 (Hauser et al., 2020; Mauduit et al., 2022). Ascl3 is a basic helix-loop-helix transcription factor expressed in a select subset of salivary gland duct cells (Arany et al., 2011; Bullard et al., 2008). Initially described to be salivary gland-specific (Yoshida et al., 2001), subsequent work has revealed that Ascl3 is expressed in numerous tissues including the lacrimal glands, trachea, stomach, and in specialized microvillar cells of the olfactory epithelium (Weng et al., 2016), which were recently identified as ionocytes (Ualiyeva et al., 2024).

In this study, we have generated an estrogen receptor-inducible Cre strain (*Ascl3*^*P2A-GCE*^) that maintains endogenous expression of the *Ascl3* gene. Using this allele in combination with R26^TdT^, GCaMP6f^flox^, and R26^DTA^ mouse strains, to fluorescently label or specifically ablate Ascl3-expressing ionocytes, we investigated characteristics of the Ascl3+ ionocyte cells focusing on those present in the ducts in the murine salivary gland. The results contribute new insights into the functional role of Ascl3+ ionocytes and provide a model for future research in the salivary glands.

## Materials and Methods

### Ascl3^P2A-GCE^ strain generation and genotyping

The *Ascl3*^*P2A-GCE*^ strain (Jackson Laboratory Stock 038129) was generated on the C57BL/6 background in the Transgenic Facility at the University of Rochester. A synthetic 9.7 kb DNA fragment was generated including 5′ and 3′ untranslated regions (UTRs) and the coding region of the *Ascl3* gene locus, linked with an expression cassette (GCE). The GCE is comprised of a fusion of coding sequences for a monomeric green fluorescent protein (AcGFP; Clontech) and Cre recombinase (Cre) and the tamoxifen-inducible estrogen receptor (ERT2) (Mugford et al., 2008). Prior to synthesis, the CreERT2 coding sequence was manually codon-optimized (Tang et al., 2019). The P2A-GCE cassette was inserted into the *NheI* site downstream of the Ascl3 termination codon (Fig. S1A). The neomycin (Neo) selection gene, flanked by flippase recombinase target sites (FRT), was inserted following the GCE cassette. The targeting construct was linearized with Sac*II* and Asc*I* and electroporated into C57BL/6J-129S6 hybrid embryonic stem (ES) cells.

Homologous recombination was assessed using long-range PCR with 3 sets of primers: (1) GFP PCR: AcGFP 33-60F: 5’-GGC ATC GTG CCC ATC CTG ATC GAG CTG A-3’ and AcGFP 541-14R: 5’-GCC AGC TGC ACG CTG CCA TCC TCG ATG T-3’; 58° annealing temperature, 500 bp product; (2) 3’LR PCR: Neo5mer2: 5’-CAT GCT CCA GAC TGC CTT G-3’ and Ascl3 3’LRF: 5’-GGACAGAGACTGGTACCTGGTGAATGTGCAGT-3’; 62° annealing temperature, 4.5 kb product; (3) 5’LR PCR: Ascl3 5’LRF: 5’-CCCAGACTACCCTTGAACTTACTCTGTAGACCA-3’ and PA GCE 85-59R: 5’-CCTCCACGTCTCCAGCCTGCTTCAGCA-3’; 64° annealing temperature, 0.8 kb product (Fig. S1A).

Five positively targeted ES clones were identified and two were injected into C57BL/6J blastocysts to generate mouse chimeras. Chimeras were mated with C57BL/6J mice to generate heterozygous *Ascl3*^*P2A-GCE/+*^ mice. The Neo selection cassette was removed from the resulting progeny by crossing with the B6.129S4-*Gt(ROSA)*^*26Sortm1(FLP1)Dym*/RainJ^ (FLPe) strain (Jackson Laboratory Stock 009086) to introduce FLP recombinase. To confirm FLP-mediated Neo deletion, PCR was performed with the primers: P2A GCE 2547-2567F: 5’-CCTGACCCTGCAGCAGCAGCA-3’ and Ascl3 3′ LRF: 5’-CCAGCATGGGACTCTTGGTCTACATGAGCT-3’; 65° annealing temperature, 523 bp product. Mice positive for *Ascl3*^*P2A-GCE*^, and negative for Neo, were backcrossed with C57Bl/6 to remove the FLP allele. Generation of a 523 bp band provided evidence for removal of the Neo gene. The *Ascl3*^*P2A-GCE*^ strain is maintained as heterozygotes and backcrossed to C57Bl/6.

### Endogenous Ascl3 expression

SMGs were isolated from six-month-old females (3 C57BL/6 and 3 *Ascl3*^*P2A-GCE*^/+). RNA extraction was done using the Total RNA mini kit (Omega Biotek; R6834-01), with RNAse-free DNAse treatment Qiagen (79254). RNA quantification was measured by nanodrop. cDNA was prepared using M-MLV reverse transcriptase kit (Invitrogen; 28025013) and 5 ug of RNA from each sample. RT-PCR was done using primers to detect Ascl3 coding sequence (Ascl3 CDSF: 5’-GGTGAAAGGAAACGATGGACACC; and Ascl3 CDSR 437-417: 5’-TTCTCCAGGTAGTCCTCCGGC); and murine ribosomal protein l32 (LE32) primers as control (LE32F: 5’-TTCATCAGGCACCAGTCAGACC; and LE32R: 5’-ACACAAGCCATCTACTCATT). PCR was done with PowerPol 2X PCR Mix with Dye (Abclonal; RK20719); the PCR program was 94°/60°/72°C for 35 cycles.

### Strains generated for analysis using Ascl3^P2A-GCE^ Cre allele

To label ionocytes *in vivo* for subsequent analysis, *Ascl3*^*P2A-GCE*^/+ heterozygotes were mated with the B6.Cg-*Gt(ROSA)26Sor*^*tm9(CAG-tdTomato)Hze*^/J (R26^TdT^; Jackson Laboratory Stock 007909) reporter strain. To label ionocytes for detecting single cell action potentials, *Ascl3*^*P2A-GCE*^/+ heterozygotes were mated with homozygous B6J.Cg-*Gt(ROSA)26Sor*^*tm95*.*1(CAG-GCaMP6f)Hze*^/MwarJ (R26^GCaMP6f^; Jackson Laboratory Stock 028865). To enable cell-specific ablation of ionocytes, *Ascl3*^*P2A-GCE*^/+ heterozygotes were mated with heterozygotes of *Gt(ROSA)26Sor*^*tm1(DTA)Jpmb*^/J strain (R26^DTA^/+; Jackson Laboratory Stock 006331) [Ivanova et al., 2005]. Genotyping of *Ascl3*^*P2A-GCE*^/+; R26^TdT^, *Ascl3*^*P2A-GCE*^/+; R26^GCaMP6f^ and *Ascl3*^*P2A-GCE*^/+; R26^DTA^ heterozygotes was performed using the primers Ascl3 forward 5′-CCACCCCAGTGCCTCTACACAAAT-3′, Ascl3 reverse 5′-GTCGCTGGAGAAGGGCAGCAGA-3′ in combination with allele-specific primers recommended by the supplier (Jackson Laboratory). Investigation of ionocyte Ca^2+^ activity was also performed in Tg(KRT14-cre)^1Amc^/+; R26^GCaMP6f^/+ heterozygotes using the Tg(KRT14-cre)^1Amc^ strain (Jackson Laboratory Stock 004782). Mice were maintained on a 12-hour light/dark cycle in a one-way, pathogen-free facility at the University of Rochester Medical Center. Food and water were provided ad libitum. All procedures were approved and conducted in accordance with the University of Rochester IACUC.

### Tamoxifen administration

TdTomato expression was activated in ionocytes through tamoxifen (Sigma-Aldrich; T5648) administration to 6-week-old male and female *Ascl3*^*P2A-GCE*^/+; R26^TdT^/+ double heterozygote mice by oral gavage at 0.25 mg/g of body weight. Mice received either one or two doses of tamoxifen on consecutive days. To induce the expression of R26^GCaMP6f^, tamoxifen was given to *Ascl3*^*P2A-GCE*^/+; R26^GCaMP6f^ and to Tg(KRT14-cre)^1Amc^/ R26^GCaMP6f^ male and female adult mice by oral gavage at a dose of 0.25 mg/g of body weight for 3 consecutive days. Cell-specific ablation of ionocytes was induced by treating *Ascl3*^*P2A-GCE*^/+; R26^DTA^/+ adult male and female mice with one dose of tamoxifen at 0.25 mg/g of body weight by oral gavage. For saliva pH experiments, both *Ascl3*^*P2A-GCE*^/+ control and *Ascl3*^*P2A-GCE*^/+; R26^DTA^/+ male and female mice were given one dose of tamoxifen at 0.25 mg/g of body weight by oral gavage.

### Immunohistochemistry

Tissues were collected for analysis at 1 or 2 weeks after tamoxifen administration and fixed in 4% PFA overnight at 4°C. Tissues were processed using a Tissue-Tek VIPTM processing machine (Sakura Finetek USA) and embedded in paraffin. Blocks were cut into 5 µm sections collected on Superfrost Plus slides (Thermo Scientific). Sections were deparaffinized, then heated in 10 mM Tris base, 1 mM EDTA for 10 minutes for antigen retrieval, then blocked with 10% normal donkey serum (Jackson ImmunoResearch Labs, 017-000-121)/1% BSA/0.1% Triton X-100 in PBS for 1 hour. Alternatively, with anti-mouse antibodies MOM blocking was done for 1 hour followed by addition of antibody in MOM diluent. Sections were incubated with antibody in PBS with 1% bovine serum albumin overnight at 4°C. Primary antibodies used: rabbit anti-RFP (1:500; Rockland Immunochemicals, 600-401-379), mouse anti-RFP (1:500; Takara, 632392), rabbit anti-Nkcc1 (1:500; Cell Signaling Technology, D208R), mouse anti-IP3R3 (1:400; BD Bioscience, 610312), rabbit anti-CFTR (1:200; Cell Signaling Technology, 78335), goat anti-Nkcc1 (1:200; Santa Cruz, sc-21545) and mouse anti-Beta tubulin (1:1000; Abcam, AB78078). After 3 washes in PBS, sections were blocked in 10% NDS at room temperature 1 hour, then incubated with secondary antibodies diluted 1:500 in 0.1% BSA and incubated 1 hour at room temperature. Secondary antibodies used: donkey anti-mouse IgG Alexa Fluor 594, donkey anti-rabbit IgG and Alexa Fluor 488, donkey anti-rabbit IgG Alexa Fluor 594, donkey anti-goat IgG Alexa Fluor 488 (Thermo Scientific, A21203, A21206, A21207, A11055). After 3 PBS washes, sections were stained with DAPI (1:500, Thermo Scientific), rinsed and mounted with Immu-mount^TM^ solution (Thermo Scientific). Lacrimal glands were dissected from mice and fixed in 4% paraformaldehyde (PFA) in PBS (pH 7.4) for 1 hour. Samples were embedded in OCT, snap-frozen in 2-methylbutane (isopentane; Millipore/Sigma cat #277258, St. Louis, MO, USA) cooled with liquid nitrogen and sectioned at 10 µm thickness using a Hacker/Bright OTF5000-LS004 cryostat (Hacker Industries Inc., UK). Frozen sections were then blocked with 5% bovine serum albumin (BSA) in Tris-buffered saline containing 0.05% Tween-20 (TBST) and incubated with the indicated primary antibodies overnight at 4°C and stained with appropriate secondary antibody.

### Immunohistochemistry for STED microscope imaging

Sections were stained with either RFP and Beta-tubulin or with Nkcc1 and Beta-tubulin, as follows: sections were blocked in MOM diluent for 5 min at RT, rinsed in PBS, and then stained with mouse anti-Beta tubulin III (1:1000; Abcam ab78078) O/N at 4°C. After 3x PBS wash, secondary anti-mouse IgG Alexa 594 (1:500; Thermo Scientific, A21203) was added for 1 hour. Slides were washed 3x PBS, then blocked with 10% normal donkey serum (NDS) 1 hour at RT. After 2x PBS wash, rabbit anti-RFP (1:500 in MOM diluent; Rockland Immunochemicals, 600-401-379) or anti-Nkcc1 (rabbit; 1:200 in MOM diluent; Cell Signaling Technology, D208R) was added and incubated overnight at 4°C. After PBS wash, slides were incubated with the secondary Star Red donkey anti-rabbit IgG (1:200; Abberior, STRED-1010) RT for 1 hour, then mounted using Prolong ™ Diamond Antifade Mounting media (P36965; Invitrogen). Slides were stored at 4°C until imaged.

### Stimulated emission depletion (STED) image analysis

Stimulated Emission Depletion super resolution images were collected using Abberior Easy3D STED microscope equipped with an Olympus UPlanApo 100x/1.4NA objective lens (Expert Line-Abberior Instruments). Samples were excited with 561nm and 640nm lasers for Alexa 594 and Star Red, respectively. The 775nm laser line was used for depletion of both channels. Three-Dimension images were collected at 150nm z-step intervals. Individual Ascl3+ ionocytes (stained with anti-rabbit RFP or anti-rabbit Nkcc1) and cells labeled with mouse anti-Beta-tubulin were segmented and a separate volumetric surface was created for each ionocyte. Imaris Software was used to calculate the shortest distance between each of the labeled ionocytes and the nearest tubulin surface. As a resolution control, the extent of overlap between each ionocyte and Beta-tubulin volume was measured. The overlapping volume between the two segments were minimal indicating that the resolution was sufficient to separate the two structures. The level of resolution is estimated to be 100-150 nm or better. Additionally, the fluorophores labeling ionocytes and Tubulin were inverted to exclude wavelength-related depletion artifacts.

### In Vivo imaging

*Ascl3*^*P2A-GCE*^*/+; R26*^*GCaMP6f*^*/+* and *K14*^*Cre*^*/+; R26*^*GCaMP6f*^*/+* double heterozygous mice were used for Ca^2+^ imaging of ionocytes *in vivo*. To generate *K14*^*Cre*^*/+; R26*^*GCaMP6f*^ transgenic mice, homozygous R26^GCaMP6f^ mice (Jackson Laboratory; Jax 028865) were crossed with homozygous Tg(KRT14-cre)1Amc/J mice (Jackson Laboratory; Jax 004782). Tamoxifen (Sigma-Aldrich; T5648) was given to the mice via oral gavage at a dose of 0.25 mg/g of body weight for 3 consecutive days to excise the loxP sites flanking the STOP codon allowing expression of the cre-dependent Ca^2+^ indicator. The mice were anesthetized and the gland exposed, as described previously (Huang et al., 2024; Takano et al., 2021; Wahl et al., 2023). The immobilized gland was secured within the holder using a cover glass and maintained in Hank’s salt solution (HBSS). Ca^2+^ imaging was conducted *in vivo* via two-photon microscopy using an Olympus FVMPE-RS system equipped with an Insight X3 pulsed laser (Spectra-Physics) utilizing a heated (OKOLab COL2532) 25x water immersion lens (Olympus XLPlan N 1.05 W MP). GCaMP6f was excited at 950 nm and emission collected between 495–540 nm, with images captured at 0.5-second intervals. Imaging depths between 10 and 55 µm from the surface of the gland were routinely utilized. Photo multiplier tube settings were fixed at 600V, 1X gain, and 3% black level, with excitation laser power adjusted per animal and according to imaging depth and the field of view. Neural stimulation was accomplished through a pair of tungsten wires (WPI, Inc) inserted into the nerve bundle associated with the blood vessels entering the gland. Stimulation was generated using a stimulus isolator (Iso-flex, A.M.P.I.) set 5 mA, 200 µs, at the indicated frequency with train frequency and duration controlled by a train generator (DG2A, Warner Instruments). Statistical analyses were performed with two-way ANOVA with multiple comparisons using Prism (GraphPad) as indicated in the figure legends.

### Saliva collection following Cre-mediated ablation of Ascl3-expressing cells

Male and female adult mice at 6-to 10-months of age were used in two groups of *Ascl3*^*+*^*/+; R26*^*DTA*^ */+* control (N=6) and *Ascl3*^*P2-GCE*^ */+; R26*^*DTA*^ */+* experimental (N=8) animals. Saliva collections were performed on fasted animals at 4 or 5 days after a single dose of tamoxifen. Food was removed from the cage, and all saliva collection experiments were conducted at the same time of day (9 AM), after a 2-hour period of fasting. Water was available with no restriction. Mice were anesthetized with ketamine (75 mg/kg) and xylazine (7.5 mg/kg) and placed ventrally on a gel pad warmed to 37 degrees. Saliva secretion was stimulated by injection of pilocarpine HCl (0.01-0.5 mg/kg IP). Pre-weighed microcapillary tube was inserted into oral cavity and saliva was collected gravimetrically for 12 - 15 minutes and stored on ice. Saliva volume was calculated, and pH was measured immediately. Mice were allowed to recover and returned to cages with free access to food and water. Mice used for saliva collection were not used for tissue collection.

## Results

### Generation of inducible Ascl3^P2A-GCE^mouse strain

We had previously generated a mouse strain in which the *Ascl3* gene locus was targeted with an expression cassette for EGFP-Cre (*Ascl3*^*EGFP-Cre*^; Jackson Laboratory Stock 021794) (Bullard et al., 2008). However, the constitutive expression of Cre recombinase and complete removal of the Ascl3 coding sequence in this strain limited its use. Here we report the generation of an inducible Cre strain driven by *Ascl3*, which maintains expression of the Ascl3 mRNA. The targeting construct included the *Ascl3* promoter and complete protein coding sequence, followed by a fusion cassette encoding a monomeric green fluorescent protein (AcGFP) and Cre recombinase (Cre) fused to a tamoxifen-inducible estrogen receptor (ERT2), hereafter referred to as the GCE cassette (Fig. S1A). A peptide 2A (P2A) sequence preceded the GCE cassette. The encoded 2A peptide acts to promote ribosomal skipping (Donnelly et al., 2001; Kim et al., 2011) allowing transcription of separate proteins from a single open reading frame. The targeting construct also included an excisable Neo selection gene and 4.3 kb of *Ascl3* 3′ untranslated sequence. Maintenance of endogenous Ascl3 mRNA expression in *Ascl3*^*P2A-GCE*^/+ mice was confirmed by RT-PCR (Fig. S1B).

### Ascl3^P2-GCE^ drives inducible expression of RFP in the major salivary glands

*Ascl3*^*P2A-GCE*^/+ heterozygotes were mated with the R26^TdT^ Tomato reporter strain, to generate *Ascl3*^*P2A-GCE*^/+; R26^TdT^/+ double heterozygotes. Tamoxifen was administered at 6 weeks of age and expression was analyzed after 1 week using immunohistochemistry (IHC) on paraffin-embedded sections of tissues. Using this protocol, tamoxifen activation resulted in robust red fluorescent protein (RFP) expression in cells scattered through the duct in the SMG of both male and female mice (Fig. 1B-D). No exogenous RFP expression was detected after tamoxifen treatment of +/+; R26^TdT^/+ heterozygotes in the absence of Cre, or in *Ascl3*^*P2A-GCE*^/+; R26^TdT^/+ double heterozygotes in the absence of tamoxifen (Fig. 1A). All subsequent experiments were performed using a single administration of tamoxifen, as the number of RFP+ cells did not differ significantly with one or two doses (data not shown). However, we cannot rule out that increasing the number of injections or dose of tamoxifen would reveal additional populations of RFP+ cells.

**Figure. 1.**
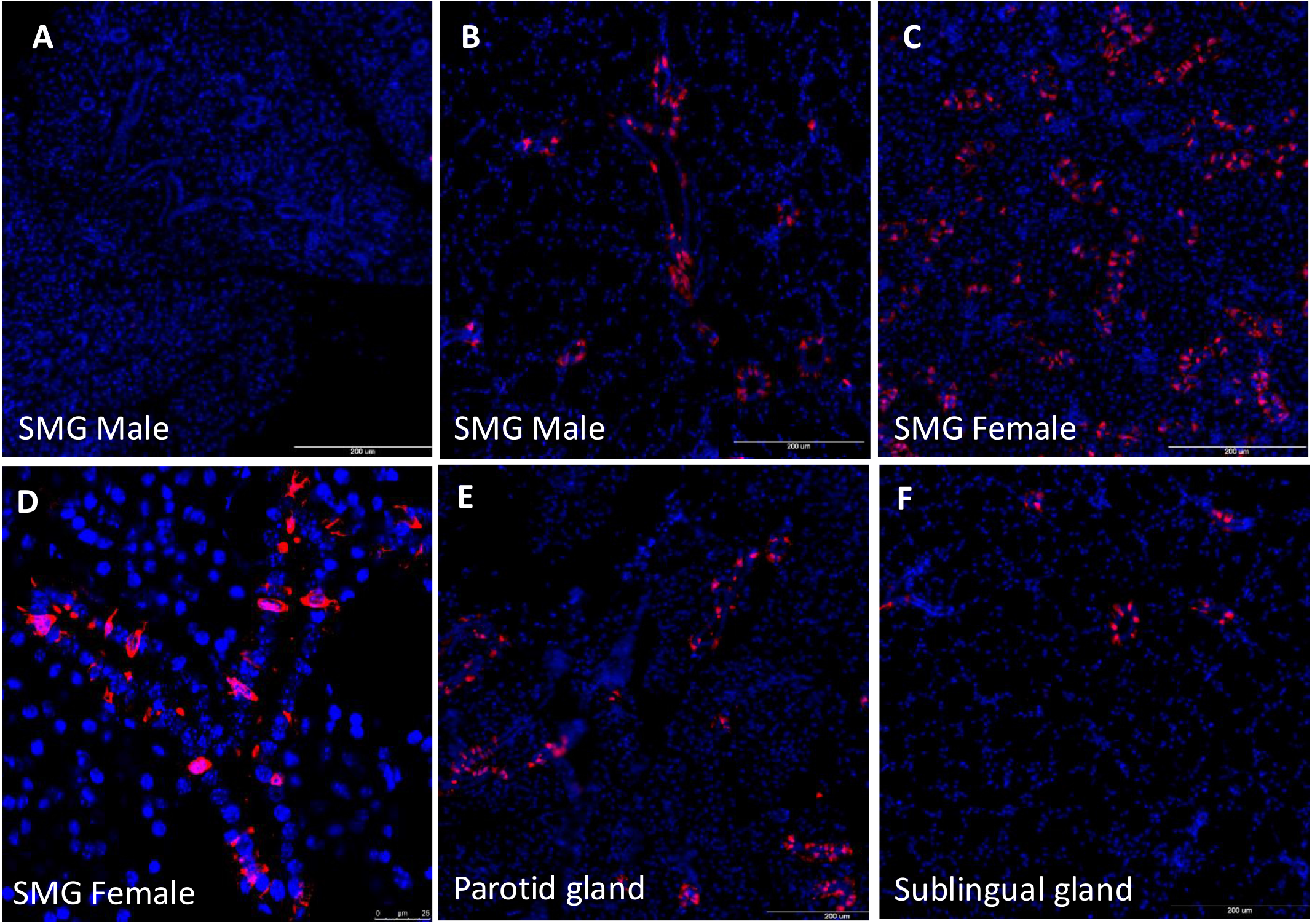
Male and female mice at 6 weeks-old were given tamoxifen on 2 subsequent days. After 2-week chase, tissues were harvested and embedded in paraffin. Sections were stained with antibody to red fluorescent protein (RFP) and DAPI. **A**. *WT +/+; R26*^*TdT*^/+ male submandibular gland (SMG), scale bar = 200 µm **B**. *Ascl3*^*P2A-GCE*^*/+; R26*^*TdT*^*/+* male SMG, scale bar= 200 µm **C**. *Ascl3*^*P2A-GCE*^*/+; R26*^*TdT*^*/+* female SMG, scale bar= 200 µm **D**. *Ascl3*^*P2A-GCE*^*/+; R26*^*TdT*^*/+* female SMG, scale bar= 25 µm. **E**. Parotid gland, and **F**. sublingual gland from *Ascl3*^*P2A-GCE*^*/+; R26*^*TdT*^*/+* female, 6 weeks-old; 1x Tamoxifen; 2-week chase. Scale bar= 200µm.

Tamoxifen administration activated RFP expression in a subset of cells resident in the duct in all three major salivary glands of *Ascl3*^*P2A-GCE*^*/+; R26*^*TdT*^*/+* double heterozygotes (Fig. 1B-F). Adult lacrimal glands had few RFP-positive cells (Fig. S1C). We have previously reported that the *Ascl3*^*EGFP-Cre*^ allele drives RFP labeling in the microvillar cells of the olfactory epithelium in mice (Weng et al., 2016), a finding which is recapitulated with the *Ascl3*^*P2A-GCE*^ strain (data not shown). Consistent with recent studies showing Ascl3 expression in mouse and human tracheal epithelia (Montoro et al., 2018; Plasschaert et al., 2018), we observed RFP expression in embryonic lung isolated from *Ascl3*^*P2A-GCE*^/+; R26^TdT^ mice at E17.5 (Fig. S1D).

### Duct cells labeled by Ascl3^P2A-GCE^/+; R26^TdT^/+ are ionocytes

In earlier work, we established that Ascl3-expressing cells in the salivary glands exhibit both molecular and morphological differences from surrounding duct cells (Arany et al., 2011; Bullard et al., 2008). scRNA studies initially performed in trachea and lungs revealed a unique cell population of ionocytes, which are characterized by expression of the transcription factors *Foxi1, Tfcp2l1* and *Ascl3* (Montoro et al., 2018; Plasschaert et al., 2018). Subsequent RNA Seq analysis in murine salivary glands identified a similar cell population and established that *Ascl3* expression specifically marks ionocytes localized to the ducts (Hauser et al., 2020; Mauduit et al., 2022). In the SMG and parotid glands, ionocytes were found to express fibroblast growth factor-10 (FGF-10), a secretory protein which may be involved in tissue regeneration, whereas FGF-10-positive cells were not detected in the ducts of the sublingual gland (Mauduit et al., 2022).

In contrast to other duct cells, Ascl3+ ionocytes express the sodium, potassium, 2-chloride co-transporter (Nkcc1) (Arany et al., 2011), otherwise localized to the basolateral membrane of acinar cells (Evans et al., 2000). Performing co-immunohistochemistry, we confirmed that the RFP+ cells in the ducts of *Ascl3*^*P2A-GCE*^*/+; R26*^*TdT*^*/+* mice co-express Nkcc1 in SMG, sublingual and parotid salivary glands (Fig. 2A-C). As previously reported (Yoshida et al., 2001), we noted fewer Ascl3+/Nkcc1+ cells in males than in females (data not shown), but the significance of this sexual dimorphism is not known.

**Figure 2.**
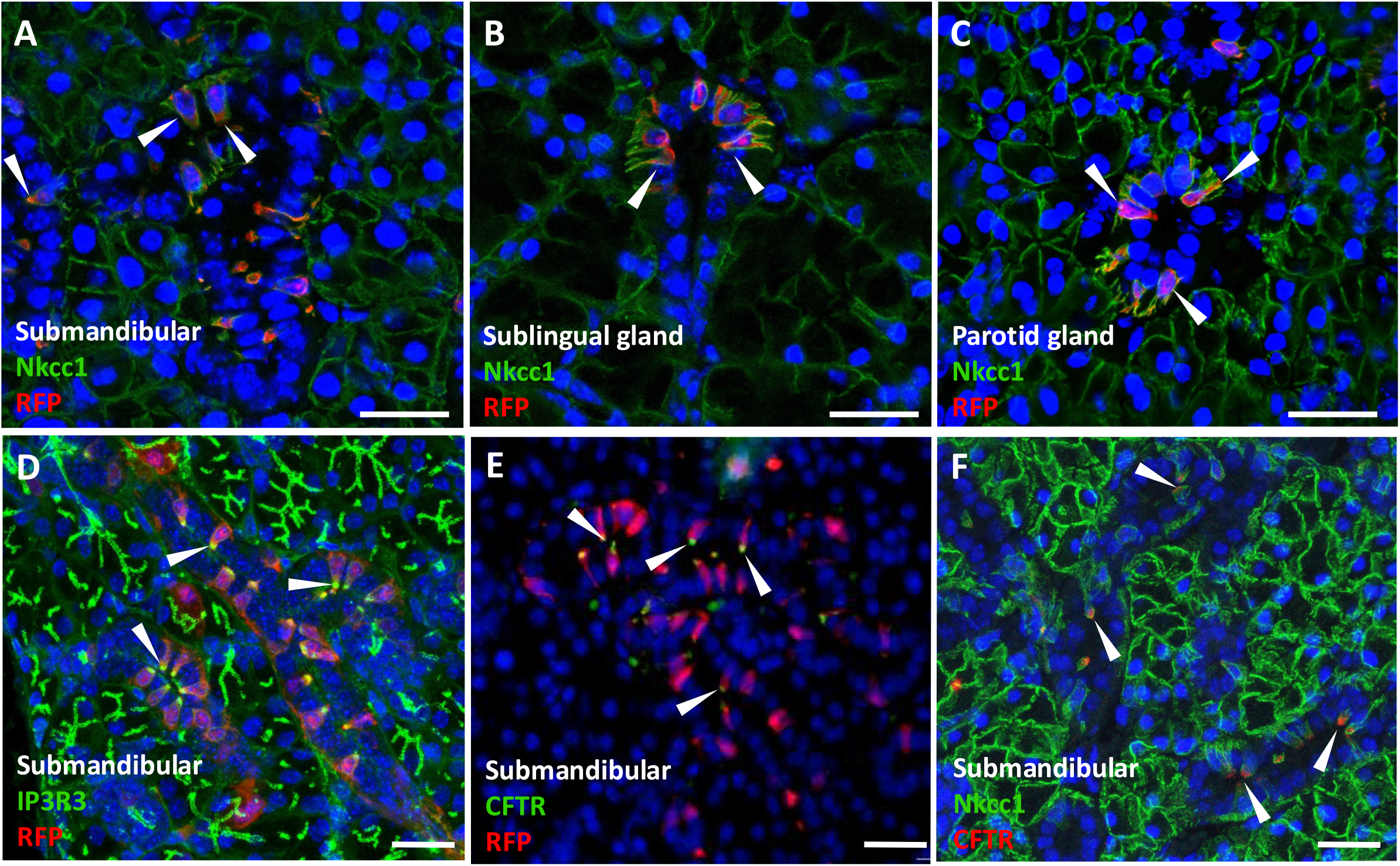
Cells labelled by RFP in *Ascl3*^*P2A-GCE*^*/+; R26*^*TdT*^*/+* mice express proteins characteristic of ionocytes. **A**. Paraffin sections of submandibular gland (SMG), **B**. sublingual gland and **C**. parotid gland, isolated from *Ascl3*^*P2A-GCE*^*/+; R26*^*TdT*^*/+* female, 6-weeks-old; 1x tamoxifen; 1-week chase; stained with antibodies to RFP and Nkcc1; nuclei stained with DAPI. White arrowheads show Ascl3+/Nkcc1+ ionocytes. **D**. Paraffin sections of SMG stained with antibodies to IP3R3 and RFP, **E**. antibodies to RFP and CFTR and **F**. antibodies to Nkcc1 and CFTR; nuclei stained with DAPI. White arrowheads indicate ionocytes expressing IP3R3+/Ascl3+, Ascl3+/CFTR+ and Nkcc1+/CFTR+. Scale bars= 25µm

In addition to Nkcc1, we found that Ascl3+ ionocytes exhibit unique expression of transmembrane proteins that are involved in acinar cell secretion, including the calcium-activated potassium channel (KCa1.1) (Arany et al., 2011) and the inositol trisphosphate receptor type 3 (IP3R3) (Weng et al., 2016) (Fig. 2D). As both IP3R3 and KCa1.1 are crucial to the secretory function of acinar cells, the expression of these proteins indicates that Ascl3+ ionocytes are involved in ion movement related to secretion.

The cystic fibrosis transmembrane conductance regulator (CFTR) is an anion selective channel exhibiting Cl− and HCO3^−^ permeability that is required for the ionic modification of primary saliva. CFTR expression was found to be a predominant feature of pulmonary ionocytes (Montoro et al., 2018; Plasschaert et al., 2018). In the murine salivary glands, CFTR expression is concentrated at the apical membrane of Ascl3+ ionocytes (Mauduit et al., 2022) (Fig. 2E,F). It has been shown that interaction between CFTR and the epithelial Na^+^ channel (ENaC) is responsible for NaCl reabsorption from the primary saliva (Catalan et al., 2010), suggesting that ionocytes modulate the ionic composition of saliva.

### Proximity of Ascl3+ ionocytes to Tubb3+ neurons

Using Nkcc1 as a marker to localize the Ascl3+ cells in SMG ducts, confocal imaging revealed flask-shaped cells with extended basal processes (Fig. 3A), like ionocytes described in pulmonary epithelia (Montoro et al., 2018), suggesting communication with surrounding cells. Chemosensory cells respond to stimulation thought to be mediated through their proximity to nerves (Xu et al., 2020). To look for innervation of Ascl3+ ionocytes in SMG, we used STED microscopy to visualize Ascl3+ cells in sections from *Ascl3*^*P2A-GCE*^*/+*; *R26*^*TdT*^*/+* mice after labeling with either anti-RFP or anti-Nkcc1 antibody in combination with antibody to Beta-tubulin (Tubb3), which is constitutively expressed in all neurons but not in glia. Imaris Software was used to calculate the shortest distance between each of the RFP+ or Nkcc1+ duct cells and the nearest Tubb3+ cell (Fig. S2). Analysis showed that 12 of 14 labeled ionocytes were located at a distance less than 100nm from the nearest Beta-tubulin-labeled cell. The two measurements indicating larger separation between ionocytes and Beta-tubulin may be explained as a sectioning artefact. Alternatively, they could represent a separate set of non-innervated cells. Most of the labelled cells were located less than 100nm from the nearest Tubb3+ surface (Fig. 3B)(Movie 1). The close proximity of the basolateral extensions to neurons in the SMG indicates that Ascl3+ ionocytes are innervated.

**Figure 3.**
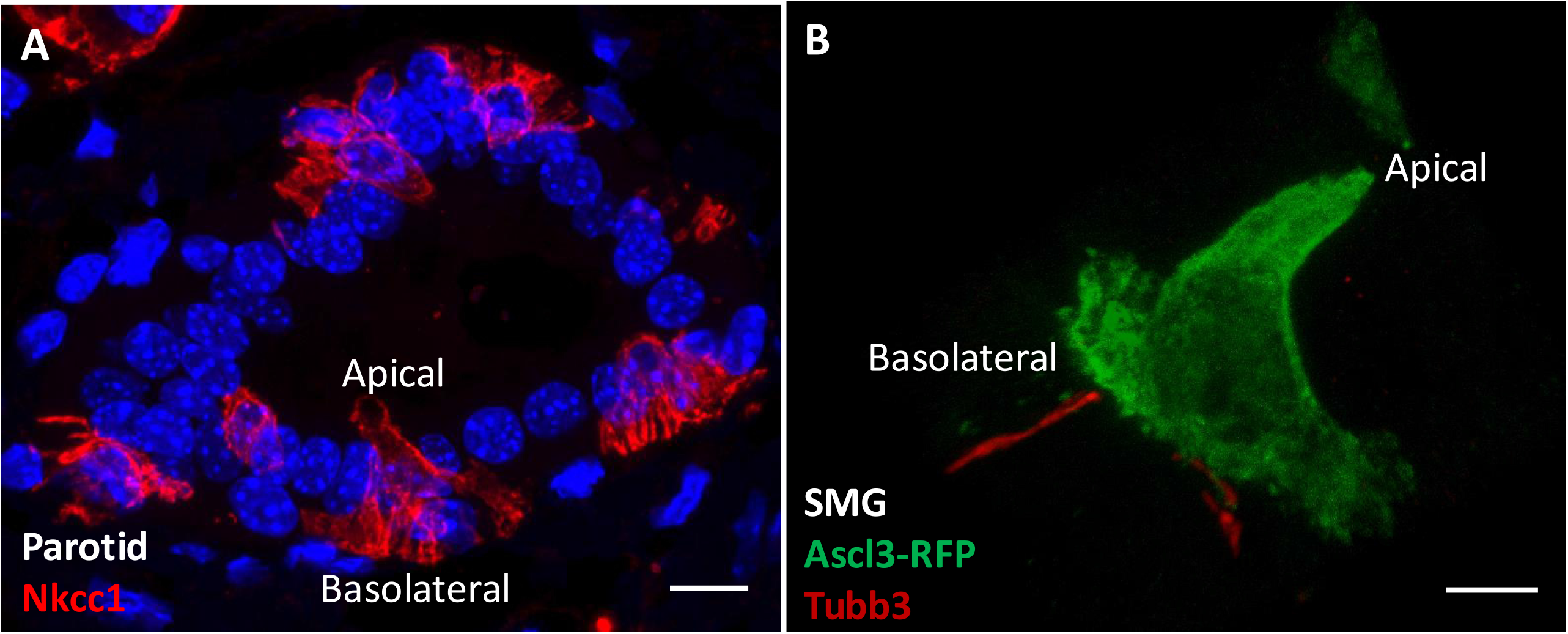
Ascl3+ ionocytes have extensive basolateral extensions adjacent to neurons. **A**. Confocal image of a section of mouse parotid gland shows duct with individual ionocytes stained using antibody to Nkcc1 (red). Nuclei are stained with DAPI. Extensive basolateral projections are visible on the cells. Scale bar= 10 µm. **B**. STED image of an RFP+ Ascl3+ ionocyte from SMG of an *Ascl3*^*P2A-GCE*^*/+; R26*^*TdT*^*/+* mouse after tamoxifen treatment. The section was co-stained using antibodies to RFP (green) and to beta tubulin (Tubb3; red). Scale bar= 4 µm.

### Cre-mediated depletion of ionocytes in SMG lowers saliva pH

To investigate ionocyte function in the SMG, we crossed the inducible *Ascl3*^*P2-GCE*^ strain to *R26*^*DTA*^. We had previously showed, using the constitutive *Ascl3*^*EGFP-Cre/+*^; *R26*^*DTA*^ strain, that the specific ablation of Ascl3+ cells in the SMG did not disrupt gland formation or the ability to regenerate after injury (Arany et al., 2011). Ascl3+ ionocytes can be detected in the SMG ducts of *Ascl3*^*P2-GCE*^*/+; R26*^*DTA*^*/+* mice due to their expression of Nkcc1 and IP3R3 (Fig. S3A). One week after a single administration of tamoxifen to induce Cre-mediated ablation through the expression of DTA, there was no detectable staining for these proteins in duct cells (Fig. S3B). Notably, the expression of IP3R3 and Nkcc1 in acinar cells was not disrupted. The ablation of Ascl3+ ionocytes was confirmed through staining for CFTR, which showed no CFTR expression in the SMG at 1 week after tamoxifen induction (Fig. 4A,B).

**Figure 4.**
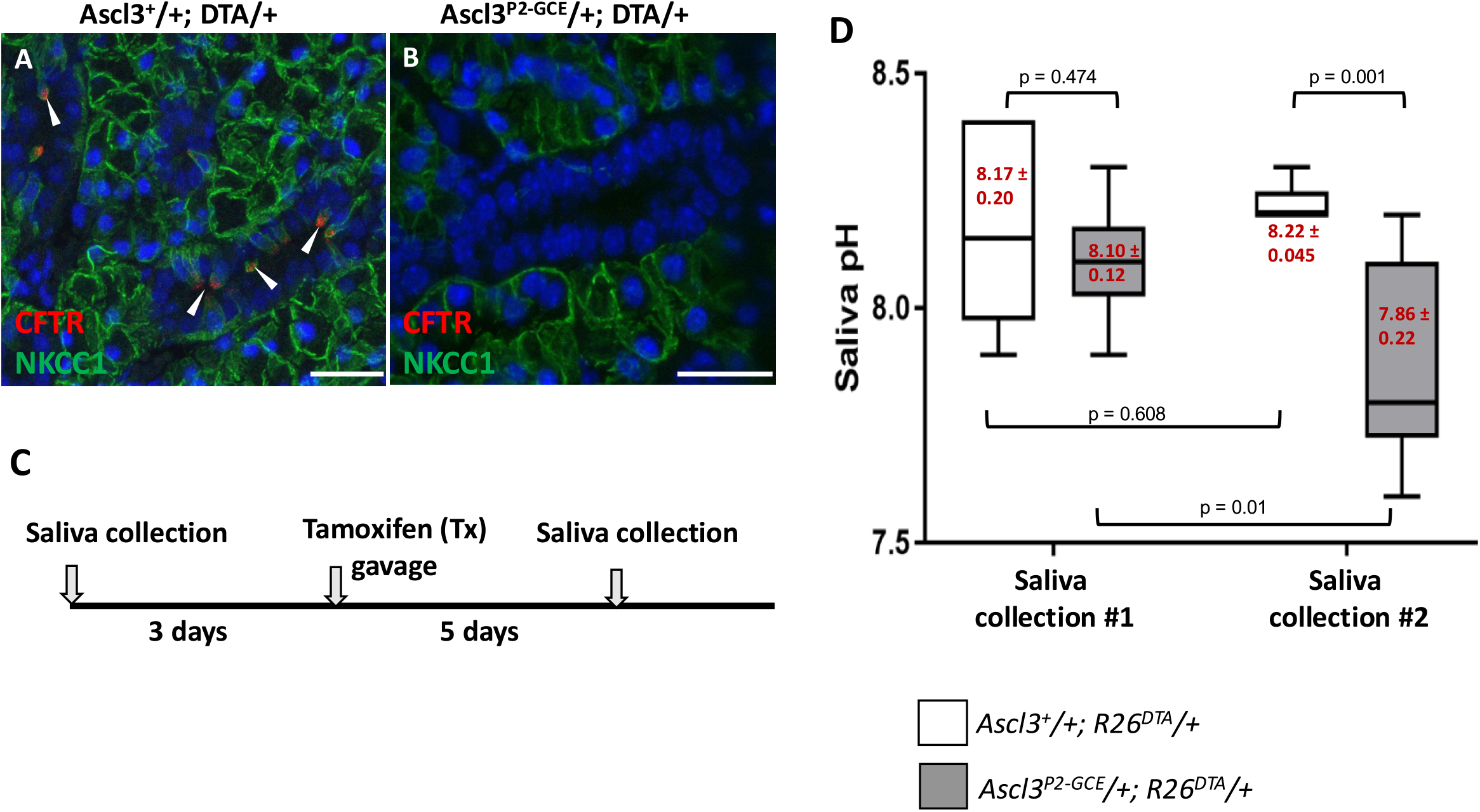
Ascl3+ cell ablation removes CFTR+ cells and lowers pH of total saliva. **A**,**B**. *Ascl3*^*P2A-GCE*^ Cre mediated ablation of Ascl3+ ionocytes removes all CFTR+ duct cells. Paraffin sections of submandibular gland (SMG) isolated from (A) Ascl3^+^; R26^DTA^ female, and (B) Ascl3^P2A-GCE^; R26^DTA^ female; 6 weeks-old; 1x tamoxifen; 1-week chase; stained with antibodies to CFTR and Nkcc1; nuclei stained with DAPI. White arrowheads in (A) show Nkcc1+/CFTR ionocytes. No detectable CFTR staining is observed following Cre-induced ablation of Ascl3+ cells in Ascl3^P2A-GCE^; R26^DTA^ mice (B). Scale bars=25 µm. **C**. Experimental timeline showing saliva collection #1, followed by administration of a single dose of tamoxifen, and subsequent saliva collection #2 at 4-5 days after Ascl3+ ionocyte cell ablation. **D**. Graph of total saliva pH measured in *Ascl3*^*+/*^*+; R26*^*DTA*^*/+* (white boxes; n = 6 (4 male, 2 female)) and *Ascl3*^*P2A-GCE*^*/+; R26*^*DTA*^*/+* (shaded boxes; n = 8 (5 male, 3 female)) before and after tamoxifen treatment and cell ablation. Both male (N=9) and female (N=5) mice were used. Data presented as mean ± SD, Two-way ANOVA, F=10.166, p=0.004

We previously reported that Ascl3+ cells exhibit a slow rate of turnover (Weng et al., 2016). However, preliminary analysis suggests that the Ascl3+ ionocytes are replenished in the SMG ducts of *Ascl3*^*P2-GCE*^*/+; R26*^*DTA*^*/+* mice within 10 days after tamoxifen administration (data not shown). Saliva collection was therefore performed on days 4 and 5 after tamoxifen treatment.

Stimulated saliva was collected from groups of adult male and female control (*Ascl3*^*P2-GCE*^*/+; +/+*) and experimental (*Ascl3*^*P2-GCE*^*/+; R26*^*DTA*^*/+*) mice, and the pH was measured (Fig. 4C). 3 days later, both control and experimental mice were administered a single dose of tamoxifen. At 4--5 days following tamoxifen treatment, stimulated saliva was again collected from each mouse and the pH was measured. The pH of total saliva collected from control animals did not change following tamoxifen treatment (8.17 ± 0.207 to 8.22 ± 0.045, p=0.608). However, the pH of total saliva collected from *Ascl3*^*P2-GCE*^*/+; R26*^*DTA*^*/+* mice after tamoxifen treatment showed a significant drop (from 8.10 ± 0.12 to 7.86 ± 0.22, p=0.004) (Fig. 4D). This indicates that ablation of Ascl3+ ionocytes disrupts the modification of primary saliva occurring in the ducts.

### Ionocytes in SMG maintain elevated intracellular [Ca^2+^] in unstimulated SMG and exhibit “blinking” activity

We crossed *Ascl3*^*P2A-GCE*^/+ mice with floxed R26^GCaMP6f^ mice to generate double heterozygous mice expressing the calcium indicator GCaMP6f specifically in Ascl3+ ionocytes. Using multiphoton microscopy (Huang et al., 2024; Takano et al., 2021; Wahl et al., 2023), we observed Ca^2+^ changes in Ascl3+ cells in SMG of live animals *in vivo*. Consistent with the localization of Ascl3+ ionocytes in striated ducts, multiphoton imaging in SMG revealed GCaMP6f fluorescence in a sparse population of cells scattered through the imaging field (Fig. 5A). Notably, no evidence of GCaMP6f expression in acinar or in non-Ascl3+ duct cells was observed. GCaMP6f fluorescence in Ascl3+ ionocytes was noticeably high under resting conditions suggesting that some Ascl3+ cells maintain elevated intracellular [Ca^2+^] ([Ca^2+^]_i_) in unstimulated SMG (Fig. 5B). Interestingly, a proportion of the bright cells underwent a rapid reduction and subsequent increase in fluorescence intensity at seemingly random frequencies in unstimulated SMG (Movie 2). We term these events “blinking”. The blinks together with the elevated basal [Ca^2+^]_i_ are a unique property of Ascl3+ ionocytes not observed in acinar or non-Ascl3+ duct cells (Takano et al., 2021; Takano and Yule, 2022; Wahl et al., 2023).

**Figure 5.**
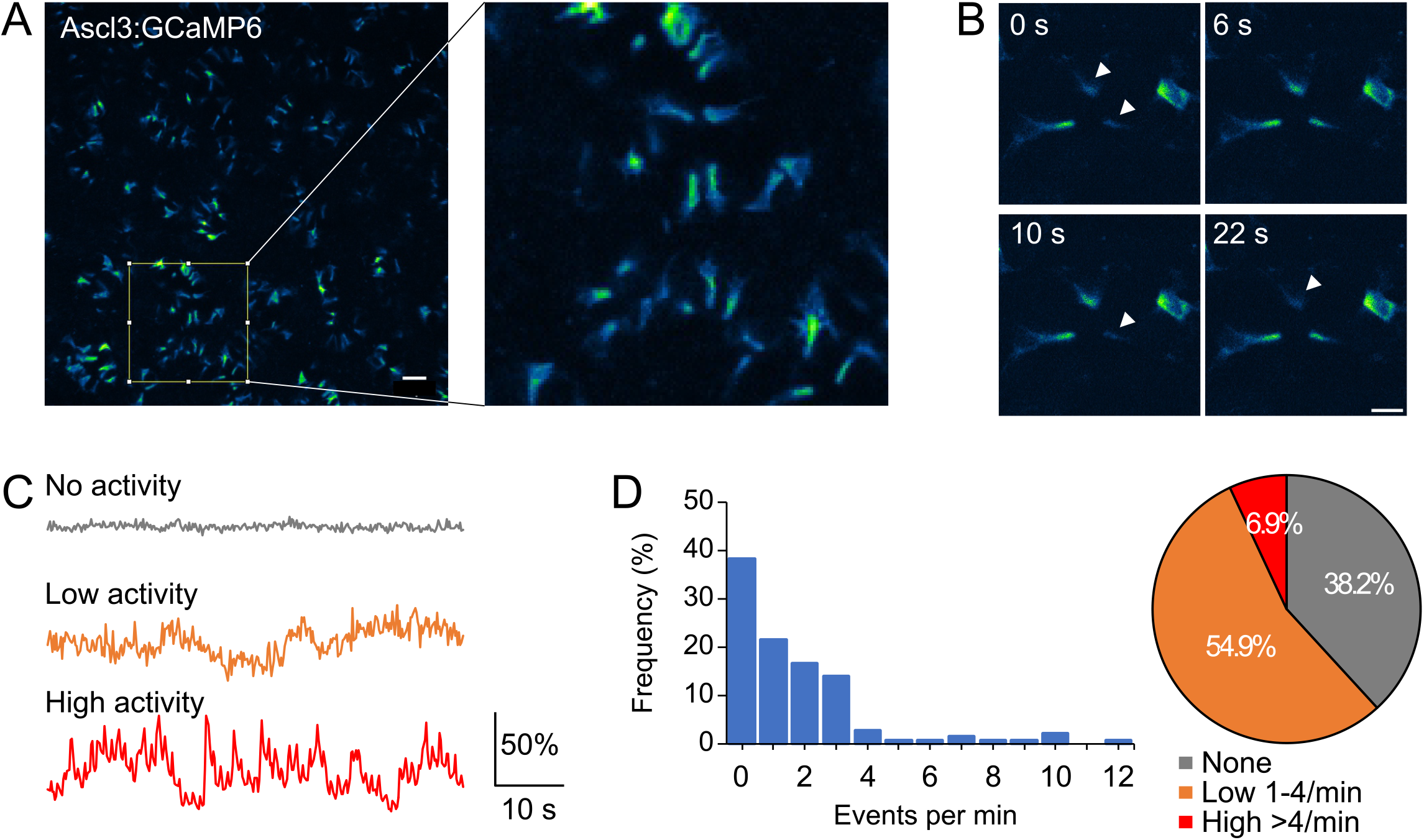
Spontaneous Ca^2+^ activity of Ascl3+ ionocytes in unstimulated SMG. **A**. Ca^2+^ imaging of submandibular gland (SMG) from *Ascl3*^*P2A-GCE*^*/+; R26*^*GCaMP6f*^/+ mouse conducted *in vivo* via two-photon microscopy. Imaging depths were between 10 and 55 µm from the surface of the gland. Scale bars= 30µm. **B**. A series of time-course images taken at 0, 6, 10 and 22 seconds depicting spontaneous blinking of Ascl3+ ionocytes in vivo. (White arrowheads indicate the cells that are transiently blinking.) 38.2% of Ascl3+ cells maintain elevated intracellular ([Ca^2+^]_I_ with no blinking in unstimulated SMG (cells with asterisks). Scale bar= 10 µm. **C**. Three representative kinetic plots of Ascl3+ cells show that they can exhibit no, low, or high blinking activities. **D**. Frequency distribution histograms of spontaneous blinking frequency depicting diverse degrees ranging from no (0 per min) to high (>4 per min) Ca^2+^ activities. N = 144 cells from 6 animals.

Not all Ascl3+ ionocytes exhibited spontaneous Ca^2+^ blinks. For example, 38.2% of the cells maintained elevated Ca^2+^ during the 60 s imaging period (Fig. 5C,D). 54.9% of the cells showed low frequency events (1-4 per min), while a small population of the cells (6.9%) showed repetitive events (Fig. 5D). Blinking cells appeared to be randomly distributed, with no obvious synchronization of events between neighboring cells.

### Nervous stimulation of the SMG decreased [Ca^2+^]_I_ in Ascl3+ ionocytes

Nervous stimulation of the SMG results in marked increases in [Ca^2+^]_i_ in acinar cells and intercalated duct cells which is important physiologically for triggering fluid secretion from the gland (Wahl et al., 2023). We investigated the effects of nervous stimulation on [Ca^2+^]_i_ signaling events in Ascl3+ ionocytes. Strikingly, using a protocol that stimulates fluid secretion (Huang et al., 2024; Takano et al., 2021), the fluorescence levels in Ascl3+ cells dramatically decreased within 5-20 s after initiation of the stimulation (Movie 3). The reduction in [Ca^2+^]_i_ persisted throughout the entire 90-second period of stimulation and continued even after the stimulation had ceased (Fig. 6A,B). Overall, the ΔF/F_0_ signal at rest was 190.5 ± 25.7% while after stimulation (5-10 Hz) it dropped to 48.2 ± 15.1% (p = 0.0037, paired t test, N = 5) (Fig. 6C).

**Figure 6.**
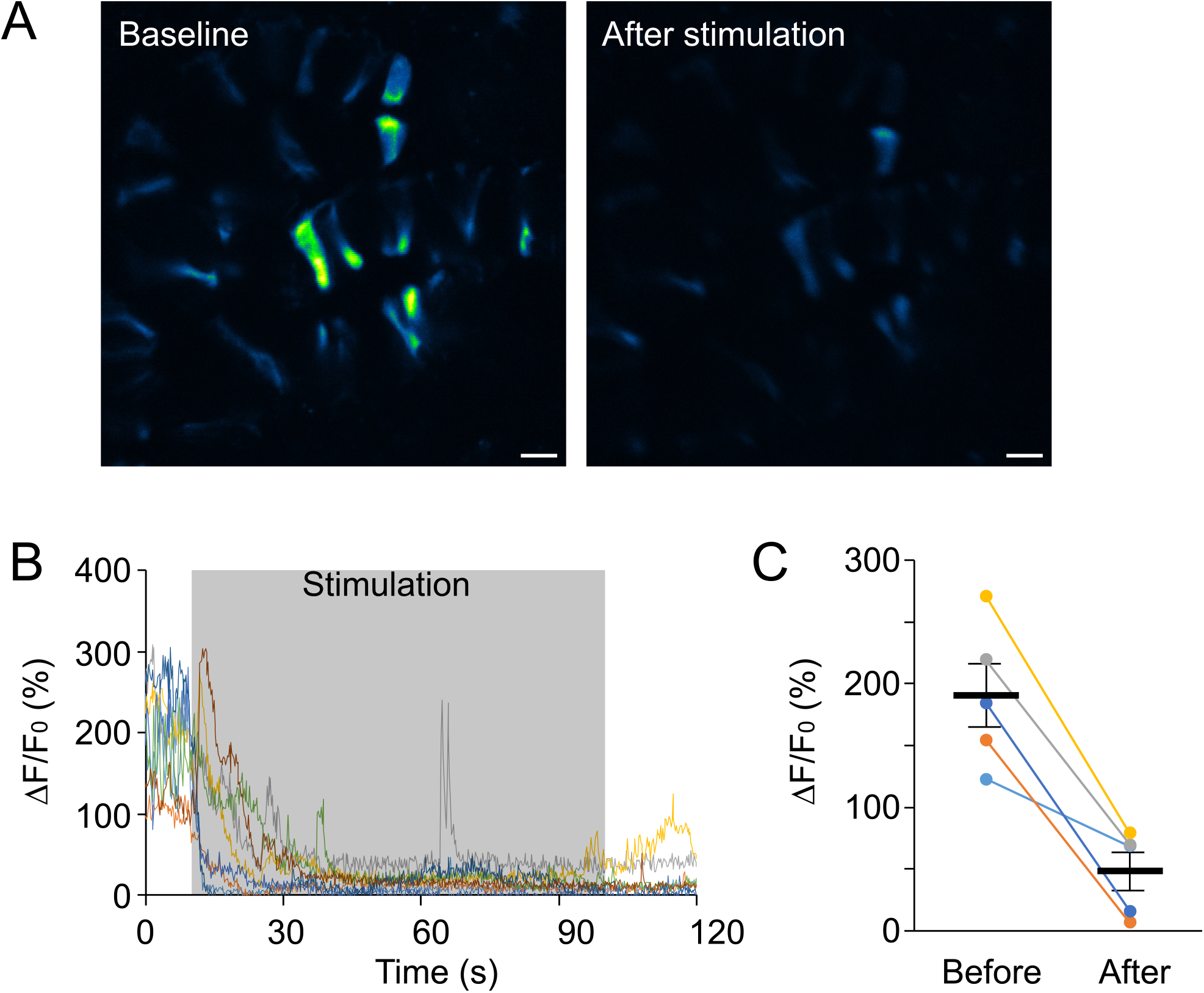
Fluorescence levels in Ascl3+ cells rapidly decrease after initiation of nerve stimulation in vivo. **A**. Average projection images of Ascl3+ cells before and 10-20 s after 10 Hz stimulation show the decrease in fluorescence. Scale bar= 10 µm. **B**. Kinetic traces of Ca^2+^ activities of individual Ascl3+ ionocytes in a field including before, during, and after a 10 Hz stimulation showing reduction in Ca^2+^ signals triggered by stimulation. **C**. A summary histogram of Ca^2+^ signals before and 10-20 s after a stimulation. Black horizontal bars indicate the averages. N = 5 animals.

Our previous *in vivo* imaging was performed in Tg(KRT14-cre)^1Amc^/+; R26^GCaMP6f^/+ mice expressing GCaMP6f driven by Keratin14 (K14) Cre (Wahl et al., 2023). As a function of K14 expression at various stages of salivary gland development, GCaMP6f is expressed in numerous distinct cell types in the SMG. In these animals we had previously noted “bright” cells scattered throughout striated ducts which we speculated were ionocytes. We investigated whether these cells had Ca^2+^ signaling characteristics consistent with Ascl3+ ionocytes. Under non-stimulated conditions, acinar and the majority of duct cells were faintly visible in the imaging field, while a sub-population of cells in the ducts exhibited very bright fluorescence (Fig. 7A). Some of the highly fluorescent cells underwent transient blinking with various frequencies in a similar fashion to Ascl3+ ionocytes. For example, 47.7% underwent transient blinking at 1-4 events per min, and 12.5% at higher frequencies (Fig. 7B,C). Similar to the observation in *Ascl3*^*P2A-GCE*^/+; R26^GCaMP6f^/+ animals, 40.3% of the highly fluorescent cells maintained an elevated Ca^2+^ level without dynamic changes. The remainder of acinar and duct cells showed no spontaneous Ca^2+^ signaling. Following nerve stimulation, acinar and non-Ascl3+ duct cells markedly increased fluorescent levels while the highly fluorescent cells exhibited a decrease in fluorescence within 5-20 seconds after initiation of the stimulation, which remained low throughout the duration of the stimulation (Fig. 7A,D). The degree of this decrease in ΔF/F_0_ signal following activation was comparable to that observed in Ascl3+ ionocytes. Putative ionocytes in K14 animals exhibited a 61.0 ± 13.0% decrease in fluorescence, compared to 72.6 ± 9.3% in *Ascl3*^*P2A-GCE*^ mice (p = 0.4883, t test, N = 5) (Fig. 7E,F). These results indicate that the Ascl3+ ionocytes exhibit a response to nerve stimulation that is distinctly opposite to that of acinar and non-Ascl3+ duct cells.

**Figure 7.**
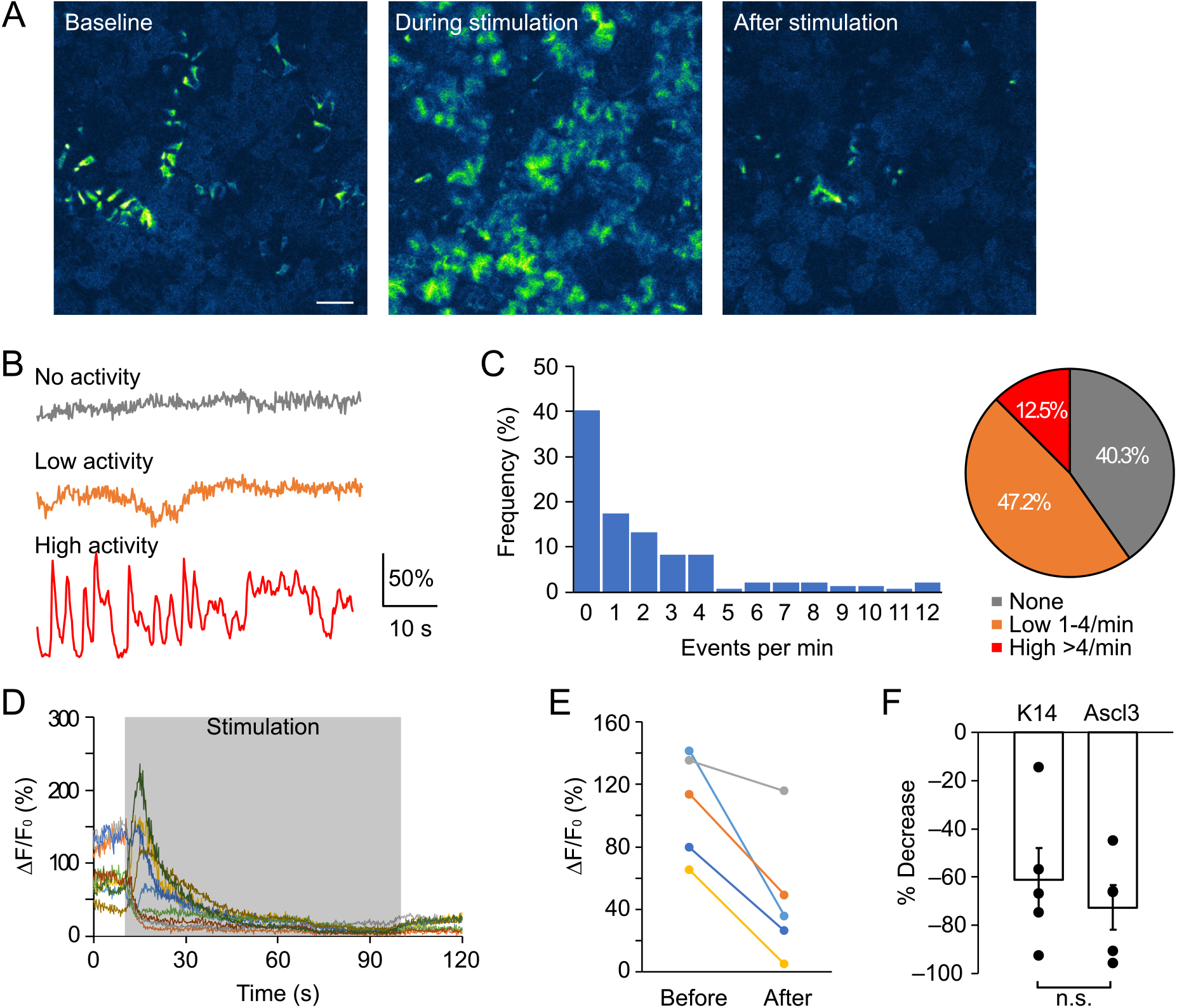
Putative ionocytes in SMG of *Tg(KRT14-cre)*^*1Amc*^*/+; R26*^*GCaMP6f*^*/+* mice show Ca^2+^ signaling consistent with Ascl3+ ionocytes. **A**. Averaged images of 10 time-course frames from SMG of *Tg(KRT14-cre)*^*1Amc*^*/+; R26*^*GCaMP6f*^*/+* animals before, during, and immediately after a 10 Hz stimulation. Scale bar= 30 µm. **B**. Three representative kinetic plots of “bright” cells in SMG of *Tg(KRT14-cre)*^*1Amc*^*/+; R26*^*GCaMP6f*^*/+* mice showing no, low, and high blinking activities prior to stimulation. **C**. Frequency distribution histograms of spontaneous blinking frequency depicting diverse degrees ranging from no (0 per min) to high (>4 per min) Ca^2+^ activities. N = 144 cells from 12 animals. **D**. Kinetic profiles showing reduction in fluorescence of putative ionocytes in SMG of *Tg(KRT14-cre)*^*1Amc*^*/+; R26*^*GCaMP6f*^*/+* mice following 10 Hz stimulation. **E**. A summary histogram of fluorescence before and after nerve stimulation. N = 5 mice. **F**. Comparison of the decrease in fluorescence seen in putative ionocytes from SMG of *Tg(KRT14-cre)*^*1Amc*^*/+; R26*^*GCaMP6f*^*/*+ mice to that observed in Ascl3+ ionocytes in *Ascl3*^*P2A-GCE*^*/+; R26*^*TdT*^*/+* mice. N = 5 mice.

## Discussion

We have previously described a subset of epithelial duct cells in murine salivary glands that uniquely express the transcription factor Ascl3 (Arany et al., 2011; Bullard et al., 2008). Recently, transcriptional profiling identified Ascl3-expressing cells as a distinct cell type known as ionocyte (Hauser et al., 2020; Huang et al., 2021; Mauduit et al., 2022). All major salivary glands have ductal Ascl3-expressing ionocytes, which exhibit distinct morphological structure and gene expression profiles. Still, the function of these cells has not been directly tested. Here, we have generated an inducible *Ascl3*^*P2A-GCE*^ Cre mouse strain and investigated additional characteristics of salivary Ascl3+ ionocytes showing that they are involved in saliva modification and display novel Ca^2+^ signaling responses.

We show that the inducible *Ascl3*^*P2A-GCE*^ strain consistently targets Ascl3+ ionocytes exhibiting unique morphological characteristics, including extended basal projectiles and polarized localization of plasma membrane channels. In contrast to surrounding duct cells, Ascl3+ ionocytes express proteins that are critical to secretory function, including Kcnma1 (Arany et al., 2011), IP3R3, Nkcc1 and Plcb2 (data not shown). Plcb2 and IP3 receptors act to stimulate calcium release from the ER intracellular stores. The Kcnma1 channel detects changes in intracellular calcium to regulate membrane potassium conductance and the membrane potential (Almassy et al., 2012). This stimulated Ca^2+^ signal regulates a range of cellular processes in salivary glands (Melvin et al., 2005). Expression of Kcnma1, IP3R3 and Plcb2 by Ascl3+ ionocytes in the salivary glands is exclusively apically localized. In contrast, Nkcc1 expression, which is normally confined to the basolateral surface of acinar cells (Evans et al., 2000), is not polarized in ionocytes. Further work is needed to understand how these differences in polarized activity relate to ionocyte function.

In addition to Ascl3, a unique feature of ionocytes in both lung and salivary gland is enriched expression of CFTR (Mauduit et al., 2022; Montoro et al., 2018; Plasschaert et al., 2018). CFTR and the epithelial sodium channel (ENaC) are co-localized at the apical membrane of pulmonary ionocytes, and are proposed to mediate Na+ and Cl− reabsorption across the luminal epithelia (Lei et al., 2023). Consistent with this, [Na+] and [Cl−] are significantly higher in saliva from Cftr knockout mice, suggesting that Cftr is involved in NaCl reabsorption in mouse SMG (Catalan et al., 2010). However, it was recently reported that pulmonary ionocytes secrete bicarbonate via CFTR-linked Cl-/bicarbonate exchange and thereby regulate pH of the airway surface liquid (ASL) (Luan et al., 2024). In agreement, the depletion of pulmonary ionocytes in ferrets impaired regulation of pH in the ASL (Yuan et al., 2023). Here, we report that specific ablation of Ascl3+ ionocytes in mouse SMG resulted in decreased saliva pH, consistent with decreased bicarbonate secretion in the absence of Cftr. Cystic fibrosis patients have mutations in the CFTR gene, and have lower saliva flow and lower saliva pH, suggesting a similar role for CFTR and ionocytes in human salivary glands (Pawlaczyk-Kamienska et al., 2019).

Neurally stimulated Ca^2+^ signaling events in acinar and intercalated duct cells are essential for saliva secretion and have been intensively studied (Takano et al., 2021; Takano and Yule, 2024; Wahl et al., 2023; Yule and Takano, 2024). However, this study is the first to describe the unique characteristics of Ca^2+^ signals in salivary gland Ascl3+ ionocytes. Unlike all other salivary gland resident cell types, Ascl3+ ionocytes exhibit high basal [Ca^2+^]_i_ levels and a subset of cells repetitively cycle between high and low [Ca^2+^]_i_ levels, an activity we term Ca^2+^ blinking. Notably, in contrast to acinar and intercalated duct cells which markedly increase [Ca^2+^]_i_ upon nervous stimulation, Ascl3+ ionocytes decrease [Ca^2+^]_i_ upon stimulation. The mechanisms underlying Ca^2+^ blinking are unclear and lead to further questions. Specifically, are Ca^2+^ blinks dependent on repetitive cycles of Ca^2+^ influx across the plasma membrane (PM) or rounds of Ca^2+^ release from the ER? In the first scenario, an oscillating membrane potential driven by the periodic activation of membrane ion channels would alter the driving force to produce periodic Ca^2+^ entry through Ca^2+^ permeable channels. Alternatively, Ca^2+^ blinks may result from Ca^2+^ release and subsequent reuptake into the endoplasmic reticulum as a result of the intrinsic biphasic activation and deactivation of the IP3R. Understanding how Ca^2+^ blinks are turned off by neural stimulation, as well as the physiological function of these signals, deserves further investigation.

Although salivary gland ducts are often considered a homogeneous population, a number of specialized duct cell types have been recognized based on structural morphology or gene expression (Mauduit et al., 2022; Sato and Miyoshi, 1997, 1998; Xiao et al., 2014). In the lung or intestine, similar specialized cell types are involved in directing immune responses to external stimuli (Branchfield et al., 2016; Gerbe et al., 2016; Howitt et al., 2016). Whether ionocytes are involved in immune responses in salivary gland ducts has yet to be determined. In addition to roles in ion transport and Ca^2+^ signaling, Ascl3+ ionocytes in the adult SMG and parotid gland also secrete FGF-10, a signaling molecule (Mauduit et al., 2022). This is consistent with our observation that Ascl3+ ionocytes are communicating with neighboring cells. While further investigation into the mechanisms is necessary, the features and activities displayed by Ascl3+ ionocytes suggest that these specialized sensory cells function as command centers in the salivary gland ducts. Based on the consistent expression of Ascl3 in ionocytes of salivary glands, lung and olfactory epithelia, the inducible *Ascl3*^*P2A-GCE*^ mouse strain provides a valuable tool for further study of this cell type, and can be utilized in a wide range of experimental setups.

## Acknowledgements

The authors thank Dr. Lin Gan for expertise in generating the inducible *Ascl3*^*P2-GCE*^ mouse strain in the Transgenic Facility at the University of Rochester, and Isabella Orup, for generating preliminary data not shown.

## Supplemental Figures

**Figure S1.**
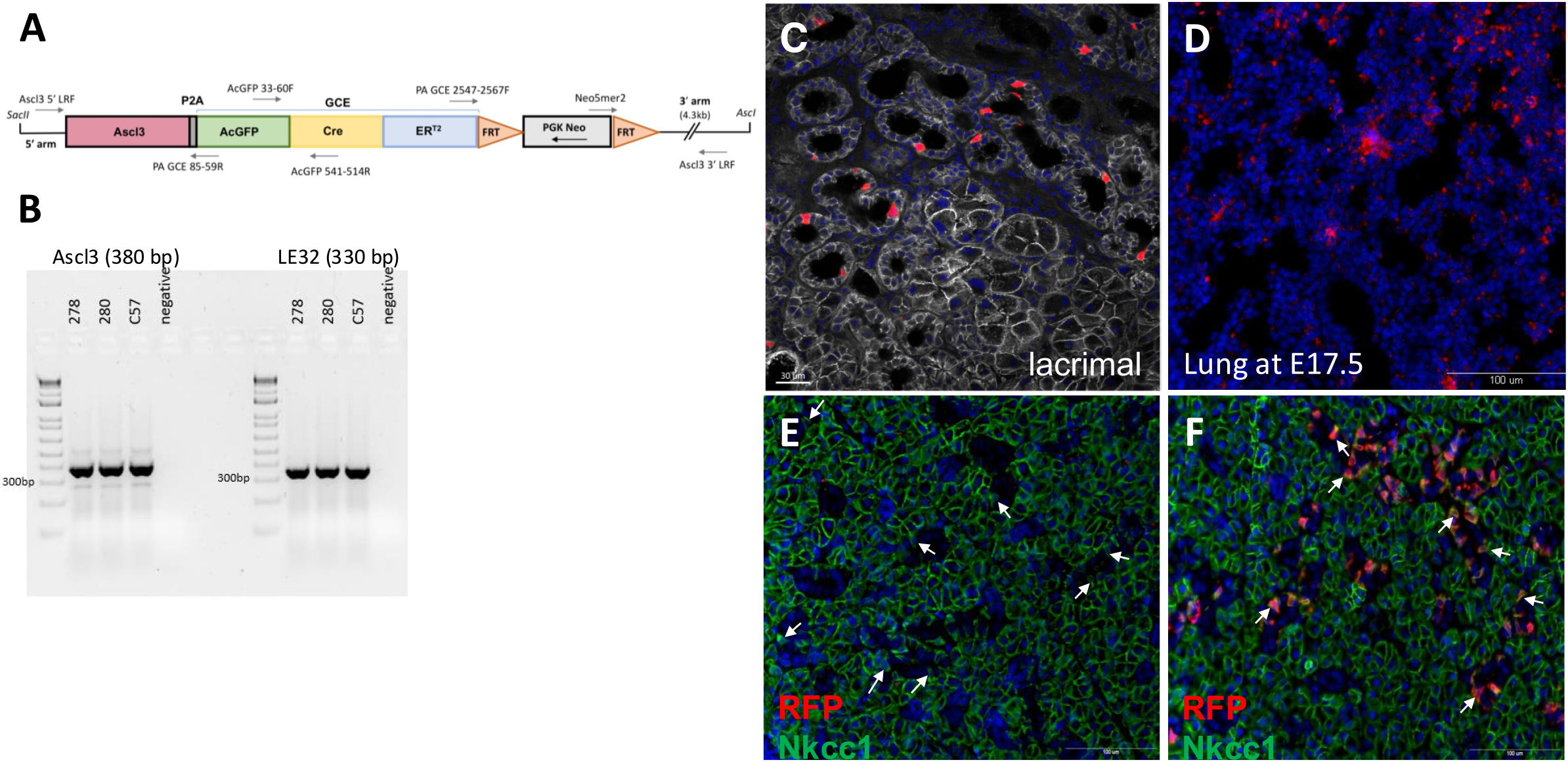
**A**. Schematic diagram of the tamoxifen-inducible Ascl3^P2-GCE^ allele, including the complete Ascl3 coding sequence, the inserted peptide 2A (P2A) sequence, and the GCE cassette comprised of the 27KDa protein derived from the jellyfish Aequorea coerulescens, (AcGFP) and the tamoxifen-inducible Cre-ER^T2^ fusion protein [GCE]. **B**, mRNA was isolated from SMG of Ascl3^P2-GCE^ and C57BL/6 female mice and used to perform RT-PCR with primers for the Ascl3 transcription factor and for the ribosomal protein LE32 for control. Lane 1, marker; lane 2, #278 Ascl3^P2-GCE^; lane 3, #280 Ascl3^P2-GCE^; lane 4, C57BL/6; lane 5, no cDNA control. Expected product for Ascl3 CDS F and Ascl3 437-417R is 380 bp; for LE32 is 330bp. **C**. Lacrimal gland isolated from Ascl3^P2A-GCE^/R26^TdT^ female at 2 weeks after 1x Tamoxifen. Frozen sections were immunostained with E-Cadherin antibody (grey; 1:200), and nuclei were counterstained with DAPI.. **D**. Embryonic lung isolated from Ascl3^P2A-GCE^; R26^TdT^ mouse embryo at E17.5 following 1x Tamoxifen given at E15.5; stained with antibody to RFP and DAPI. **E**,**F**. To confirm that activation of RFP expression is dependent on Cre-mediated recombination, Ascl3^P2A-GCE^/+; R26^TdT^/+ female was not given tamoxifen (E), while Ascl3^P2A-GCE^/+; R26^TdT^/+ female was treated at 6-weeks-old with 1X tamoxifen (F). SMGs were isolated after 1 week, fixed in 4% PFA, embedded in paraffin, and sectioned. Sections were stained with antibodies to RFP and Nkcc1. No RFP expression was observed in the absence of tamoxifen treatment. White arrows indicate Nkcc1+ duct cells previously shown to co-express Ascl3 (Bullard et al., 2008). Scale bars = 100µm

**Figure S2.**
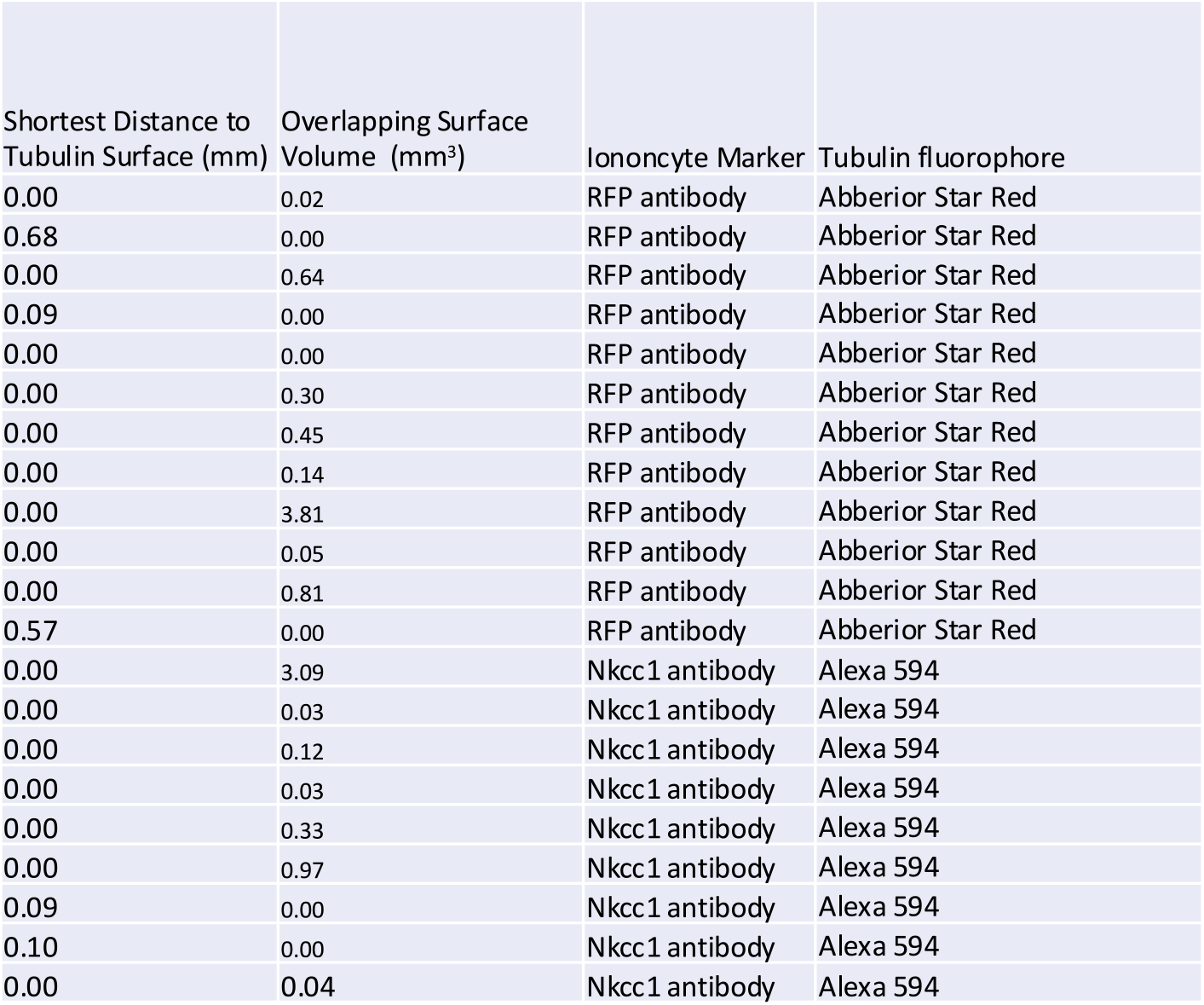
STED Image measurements of distance between Ascl3+ ionocyte and nearest Beta-tubulin 3+ neuron. Individual surfaces were created in Imaris corresponding with the signal from each Ascl3+ cell (stained with anti-RFP or anti-Nkcc1 antibody), and any neurons or neuronal segments (stained with anti-Tubb3 antibody) in every Image. Imaris Software was used to calculate the shortest distance between each of the ionocyte surfaces and the nearest tubulin+ surface, as indicated in the “Shortest Distance to Tubulin Surface” column. In the same data sets the amount of signal overlap between the ionocytes and tubulin signals (surfaces) was also measured as indicated in “Overlapping Surfaces Volume” column. The small size of the overlapping volume indicates that the two signals are separated within the resolution of the images, controlling for artifactual proximity measurements due to insufficient resolving power. Resolution in these images is estimated to be 100-150 nm.

**Figure S3.**
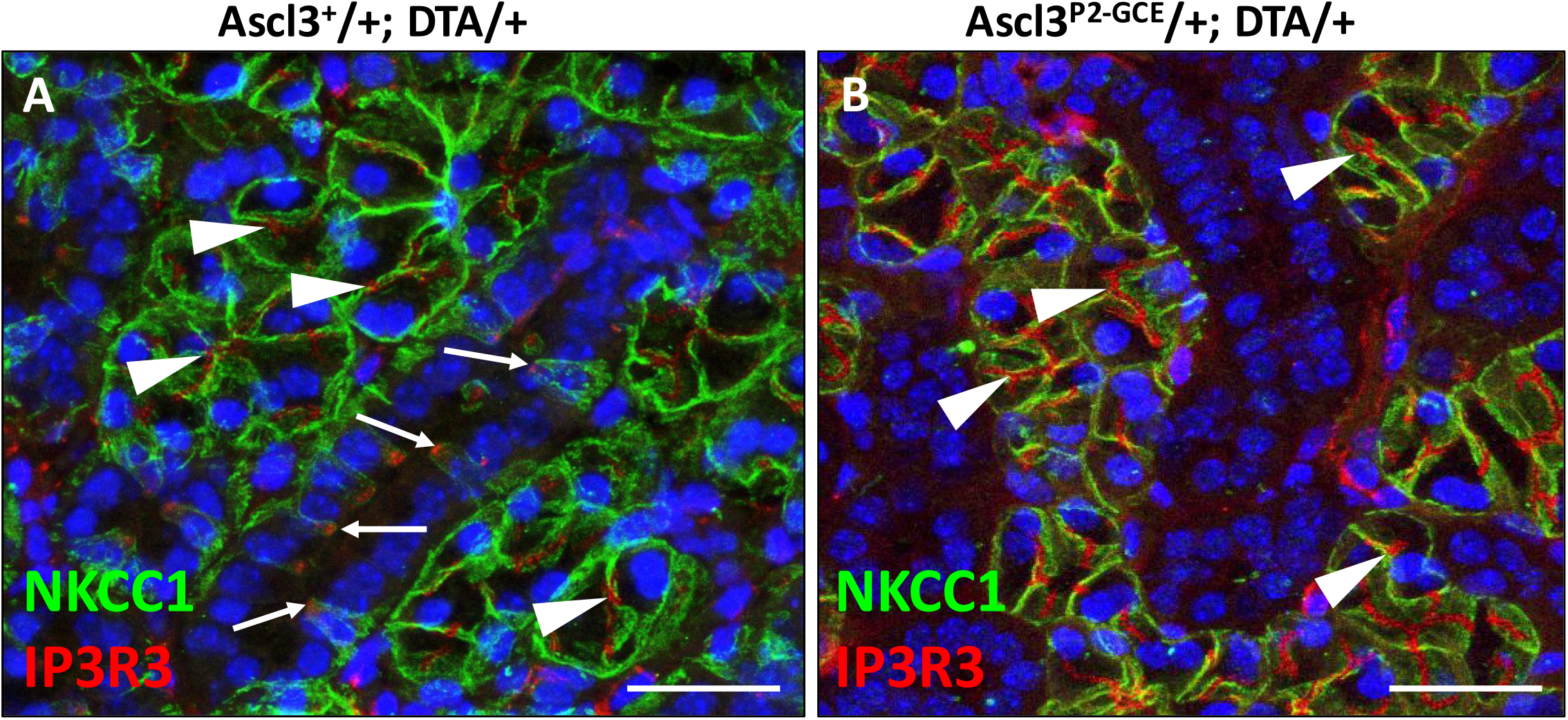
Paraffin sections of submandibular gland (SMG) isolated from **A**, *Ascl3*^*+*^*/+; R26*^*DTA*^*/+* female, and **B**, *Ascl3*^*P2A-GCE*^*/+; R26*^*DTA*^*/+* female; 6 weeks-old; 1x Tamoxifen; 1-week chase; stained with antibodies to Nkcc1 and IP3R3; nuclei stained with DAPI. White arrows in (A) indicate Nkcc1+/IP3R3+ ionocytes. IP3R3 or Nkcc1 staining is not detected in duct cells following Cre-induced ablation of Ascl3+ cells in *Ascl3*^*P2A-GCE*^*/+; R26*^*DTA*^*/+* mice (B), although expression of these proteins in the acinar cells is unaffected (white arrowheads in A and B). Scale bars= 25 µm

**Movie 1**. STED image of a duct cross-section isolated from *Ascl3*^*P2A-GCE*^*/+; R26*^*TdT*^*/+* female SMG. Ascl3+ ionocytes (green, stained with anti-RFP) and Tubb3+ neurons (red, stained with anti-Tubb3) are located in close apposition in the ducts of the murine salivary gland. The individual cell markers are distinct and well resolved spatially within the STED image. Rendered in 3D with Imaris software. Estimated resolution is 100-150 nm. Scale bar= 15 µm.

**Movie 2**. A pseudo-colored representative movie of spontaneous Ca^2+^ activity of Ascl3+ ionocytes in submandibular gland. Ca^2+^ flickering in the absence of external stimulation was observed. Frame rate, 5 Hz. Scale bar= 20 µm.

**Movie 3**. Reduction of Ca^2+^ signals in Ascl3+ ionocytes from submandibular gland induced by 10 Hz stimulation given between 10 and 100 s. Ca^2+^ reduction was observed during the stimulation period and remained decreased afterwards. Frame rate, 5 Hz. Scale bar= 20 µm.

